# The response to the DNA damaging agent methyl methanesulfonate in a fungal plant pathogen

**DOI:** 10.1101/527218

**Authors:** Shira Milo-Cochavi, Manish Pareek, Gregory Delulio, Yael Almog, Gautam Anand, Li-Jun Ma, Shay Covo

**Affiliations:** Department of Plant Pathology and Microbiology, Hebrew University, Rehovot Israel 7610001; Department of Biochemistry and Molecular Biology, University of Massachusetts, Amherst, MA USA 01003

## Abstract

DNA damage can cause mutations that in fungal plant pathogens lead to hypervirulence and resistance to pesticides. Almost nothing is known about the response of these fungi to DNA damage. We performed transcriptomic and phosphoproteomic analyses of *Fusarium oxysporum* exposed to a DNA alkylating agent (MMS). At the RNA level we observe massive induction of DNA repair pathways; the most surprising is the global genome nucleotide excision repair. Cul3, Cul4, several Ubiquitin-like ligases and other components of the protein degradation machinery are significantly induced. In agreement, we observed drug synergism between a proteasome inhibitor and MMS. While our data suggest that Yap1 and Xbp1 networks are similarly activated in response to damage in yeast and *Fusarium* we were able to observe *Fusarium* specific MMS-responsive modules. These include transcription/splicing modules that are upregulated and respiration that is down-regulated. In agreement, MMS treated cells are much more sensitive to a respiration inhibitor. At the phosphoproteomic level, Adenylate cyclase, which generates cAMP, is phosphorylated by MMS and forms a network of phosphorylated proteins that include cell cycle regulators and several MAPK pathways. Recently, there are many attempts to re-sequence isolates of fungal plant pathogens. These attempts often reveal changes in the genomes that may be explained by DNA damage exposure. Our analysis provides an important starting point in understanding how these genomic changes occur.

## 1. Introduction

In most organisms, genetic information is stored in the DNA macromolecule. DNA is subjected to many types of lesions that alter the chemical structure of either the nucleic acid bases or the sugar phosphate backbone (E.C FRIEDBERG, 2006). Lesions in DNA can originate from internal and external sources; the sunlight causes UV pyrimidine dimers, respiration produces reactive oxygen species (ROS) that cause oxidation of bases and abasic sites, and the common metabolite S-adenoysl methionine causes methylations on DNA bases (Lindahl, 1993; Lindahl and Wood, 1999; Lindahl and Barnes, 2000; E.C FRIEDBERG, 2006). Fungal plant pathogens are exposed to these types of DNA damaging agents and more. One of the best characterized responses of plants to fungal infection is the ROS burst. Within plants’ arsenal of chemical agents against pests are compounds that disrupt chromosome functions and therefore cause DNA damage, such as the lignan podophyllotoxin (Yu et al., 2017). In addition, species within microbial communities generate DNA damaging substances, which are best manifested by the aflatoxin B1 produced by *Aspergillus* species (Sengstag et al., 1996; Bullerman, 2003). DNA damage is a threat to cell survival because the modified bases often interrupt the replication machinery, thus stopping cell division. DNA damage can also block transcription, and if not repaired efficiently, severely reduces expression of essential genes. DNA damage affects not only survival but also genome integrity; it is a major determinant in mutagenesis. DNA damage can also promote structural variations in chromosomes (Argueso et al., 2008; Roberts et al., 2012; Serero et al., 2014; Ling et al., 2018). Genomic plasticity, that is often associated with low DNA repair capacity, is frequently observed in fungal plant pathogens (De Jonge et al., 2013; Jones et al., 2014; Shahi et al., 2016; Vlaardingerbroek et al., 2016). Therefore, understanding the mechanisms by which fungal plant pathogens respond to DNA damage is important.

The response to DNA damage consists of several layers, including damage recognition, signal transduction, cell cycle arrest and DNA repair. Methyl methanesulfonate (MMS) is a methylating DNA agent that causes methylation primarily on N7 of deoxyguanosine and N3 of deoxyadenosine (Beranek, 1990). Two main mechanisms repair MMS damage; the first is Mgt1, a direct reversal repair protein that removes the methylting group from the damaged base and transfers it to a cysteine residue (Lindahl and Wood, 1999; Lindahl and Barnes, 2000; Pegg, 2000). It is a suicidal enzyme in which, upon activation, its expression is induced via a positive feedback loop. The second MMS repair mechanism is base excision repair; the damaged based is removed by the Mag1 glycosylase and the remaining abasic site is excised from DNA, leaving a gap that is eventually filled and ligated (Lindahl and Wood, 1999; Lindahl and Barnes, 2000). Nucleotide excision repair machinery recognizes DNA helix distortion and is not specific to lesions; some evidence suggests that it can also be involved in MMS repair to some extent (Huang et al., 1994; Memisoglu and Samson, 2000). Homologous recombination deficient cells are very sensitive to MMS exposure, although MMS exposure does not cause breaks directly (Lindahl et al., 1997; Ma et al., 2009; Ma et al., 2011). MMS lesions that escape repair before DNA replication initiation may stall DNA replication (Harper and Elledge, 2007; Hedglin and Benkovic, 2015). The post-replication repair machinery that is governed by the Rad6-Rad18 ubiquitin conjugation enzymes allows the replication machinery to progress even in the presence of lesions (Harper and Elledge, 2007; Bansbach and Cortez, 2011; Hedglin and Benkovic, 2015; Cipolla et al., 2016; Gao et al., 2017). This could be done in an error-free or error-prone way. Error-prone post replication repair is conducted by specialized DNA polymerases (Livneh, 2001; Kunkel et al., 2003). In bacteria, a mutagenic response to DNA is best characterized in the SOS pathway. Nowadays, the notion that under genotoxic and other stresses mutagenesis programs are activated to generate genetic diversity is widely accepted (Galhardo et al., 2007). To some extent, this was also shown in eukaryotes (Shor et al., 2013). Whether or not a mutagenesis program is operated in fungal plant pathogens under stress is not known.

Breaks and gaps that occur due to MMS lesion-processing activate the DNA damage response that consists of the two major kinases ATM and ATR, which further transduce the signal to the Chk2 and Chk1 kinases (Desany et al., 1998; Ciccia and Elledge, 2010). This cascade operates cell cycle arrest that delays mitosis to provide opportunity to DNA repair mechanisms to remove lesions, thus preventing chromosome catastrophe or chromosomal aberrations during mitosis (Elledge, 1998; Harper and Elledge, 2007).

The DNA damage response was studied in high resolution in *S. cerevisiae* and *Schizosaccharomyces pombe*. These studies revealed the key DNA repair and cell cycle regulators induced at the transcription level and the post-translational modification levels (Smolka et al., 2007; Fu et al., 2008; Shalem et al., 2008; Willis et al., 2016; Zhou et al., 2016). While early observations focused on phosphorylation, more recently it became clear that ubiquitination, sumoylation, and other small protein modifications are very important (Papouli et al., 2005; Branzei et al., 2006; Moss et al., 2010; Ulrich and Walden, 2010; Hedglin and Benkovic, 2015; Cipolla et al., 2016; An et al., 2017). A systematic approach to study DNA damage response revealed activation of multiple networks (Workman et al., 2006; Bandyopadhyay et al., 2010; Guenole et al., 2013). Moreover, hundreds of proteins change their amounts or cellular localization, and their relevance to the DNA repair is unclear. Importantly, MMS that is used in some of these studies may affect other macromolecules in the cell and not only DNA (Tkach et al., 2012). To what extent the activation of all these networks in response to DNA damage is evolutionary conserved is unknown.

While key players in DNA repair in filamentous fungi were previously identified, very little is known about the entire cellular response to DNA damage (Goldman et al., 2002; Goldman and Kafer, 2004; Inoue, 2011). Almost nothing is known for fungal plant pathogens. Here we describe the cellular response of germinating conidia of *F. oxysporum* to MMS at the transcriptome and phosphoproteome level. MMS is a model compound for internal metabolites that can methylate DNA and other compounds such as the common compound S-Adenoysl methionine. Because MMS is administrated chronically it is a model for exposure of plant compounds that may damage the DNA or interfere with its replication. We observe several evolutionary conserved pathways that are activated in yeast and *Fusarium* in response to MMS exposure such as the Chk1-Chk2 pathway, the proteasome pathway, and Xbp1 Yap1 network. But we could also identified modules that are affected by MMS in *Fusarium* and not in yeast like transcription, Cul4-Nedd8 and respiration.

## 2. Materials and Methods

### 2.1 Media and growth conditions

*Fusarium oxysporum* f.sp. *lycopersici* 4287 cultures were grown in sterile KNO_3_ media (1.36 gr/l yeast nitrogen base, 24 gr/l sucrose, 100 mM KNO_3_) and incubated at 28°C with shaking at 250 rpm for five days. The cultures were filtered to obtain conidia that were washed in autoclaved ddH_2_0. Half-billion conidia were then inoculated in 100 ml potato dextrose broth (PDB, BD, Sparks USA) for five hours, then methyl methanesulfonate (Sigma) was added to a final concentration of 0.1% for another three hours while incubating at 28°C with shaking at 250 rpm. As a control, conidia were further grown for another three hours. Next, the conidia were harvested and frozen in liquid nitrogen until RNA or protein purifications.

### 2.2 RNA purification

Cells were disrupted using Minilys bead beater (Bertin Instruments) for 30 seconds at medium speed. RNA was purified using the RNAeasy Plant Mini Kit of QUIAGEN (Hilden, Germany) and was treated on-column with RQ1 RNase-free DNase (Promega Corp., USA) to remove additional residues of genomic DNA. RNA quality was measured by a tape station machine and kit (Santa Clara, USA). RNA was submitted to the Crown Institute of Genomics of the G-INCPM (Weizmann Institute of Science, Israel), and RNA sequencing libraries were made using TrueSeq Kit according to the instructions (Illumina, San Diego USA). Next, the libraries were sequenced on one lane of HiSeq 2500 machine. For lower MMS doses (0.05%, 0.01%) RNA library were made using the 3’ quantseq kit of Lexogen, followed by sequencing using the Nextseq machine of Illumina.

### 2.3 Sequencing and initial analysis

Reads were aligned to the *F. oxyspurum* 4287 (FO2) genome (downloaded on 26/Aug/2015 from the Broad Institute Fusarium comparative genome project) using TopHat (v2.0.10) (Kim et al., 2013). HTSeq-count (version 0.6.1p1) (Anders et al., 2015) was used to count reads on genes. Differential expression analysis was performed using DESeq2 (1.6.3) (Love et al., 2014) with betaPrior set to False.

### 2.4 Annotation of *Fusarium oxysporum* genes

*Fusarium oxysporum* f. sp. *Lycopersici* 4287 v2 protein sequences were obtained from JGI MycoCosm (http://genome.jgi.doe.gov/Fusox2/Fusox2.home.html) and annotated using the blast2GO platform (Conesa et al., 2005). The longest protein sequence of each gene was used as query in a BLAST search against both the NCBI non-redundant protein database (nr) (O’Leary et al., 2016), and separately, against the Swiss-Prot protein database (Bairoch and Apweiler, 2000). BLAST hits for genes were mapped to associated GO terms, and the associated GO terms were evaluated and assigned to *F. oxysporum* genes using Blast2GO. Alternatively, Functional enrichment analysis was performed using g:Profiler using Fisher’s one-tailed test. The g:SCS method was used for computing multiple testing correction with P value less than 0.05 as the significance threshold (http://biit.cs.ut.ee/gprofiler).

### 2.5 Protein extraction

The conidia pellets were re-suspended in freshly made 8M urea lysis buffer (6.006 gm urea in 12.5 ml 0.1M Tris pH 7.9) The urea buffer was supplemented with phosphatase inhibitor (Sigma, P0044)). Cells were then disrupted by Minilys bead beater for 30 seconds.

#### Sample preparation for mass spectroscopy

All chemicals are from Sigma Aldrich, unless stated otherwise. Lysate was subjected to in-solution tryptic digestion as follows: lysate was centrifuged at 16,000 g for 10 minutes to pellet cell debris, and clear lysate was transferred to a fresh tube. Proteins were reduced by addition of DTT (5mM final concentration, 50 min at 50°C) and alkylated with iodoacetamide (10 mM final concentration, 20 min at room temp in the dark). The samples were diluted 1:4 using 50 mM ammonium bicarbonate and digested with trypsin (1:50 enzyme:protein ratio) overnight at 37°C, followed by a second trypsin digestion for 4 hours at 37°C. Digestion was stopped by addition of trifloroacetic acid to 1% final concentration, desalted using Oasis HLB in μElution format (Waters, Milford, MA, USA), vacuum dried and stored in −80°C until analysis.

For phosphoproteomics, phosphopeptides were enriched from the total protein digest using a ProPac IMAC-10 column (4×50mm) (Thermo Scientific, P.N. 063276) with loading buffer (A) 0.1% TFA, 30% ACN and elution buffer (B) 0.3% NH_4_OH, pH 11.7. Peptides were loaded on the column in 10 minutes with flow 0.1ml/min (0% B). Phosphopeptides were eluted from the column using the following gradient: first elution from 10 to 15 minutes with flow 0.6 ml/min (B from 0% to 15%), second elution from 15 min to 44 minutes with flow 0.02 ml/min (B from 15% to 30%), washing from 44 minutes to 50 minutes with flow 1 ml/min (B from 30% to 50%) and following equilibration from 50 minutes to 60 minutes with flow 1 ml/min (0% B). The phosphopeptides containing peak (second elution fraction) was collected in 1.5 ml reaction vessels (~1.2 ml), speed vac. to dryness, and reconstituted in 25 μL in 97:3 acetonitrile:water + 0.1% formic acid.

### 2.6 Liquid chromatography

ULC/MS grade solvents were used for all chromatographic steps. Each sample was loaded and analyzed using split-less Nano Ultra-Performance Liquid Chromatography (10 kpsi nanoACQUITY; Waters, Milford, MA, USA). The mobile phase was: A) H_2_O + 0.1% formic acid and B) acetonitrile + 0.1% formic acid. Desalting of the samples was performed online using a Symmetry C18 reversed-phase trapping column (180 μm internal diameter, 20 mm length, 5 μm particle size; Waters). The peptides were then separated using a T3 HSS nano-column (75 μm internal diameter, 250 mm length, 1.8 μm particle size; Waters) at 0.35 μL/min. Peptides were eluted from the column into the mass spectrometer using the following gradient: 4% to 30% (B) in 155 min, 30% to 90% (B) in 5 minutes, maintained at 90% for 5 minutes and then back to initial conditions.

### 2.7 Mass Spectrometry

The nanoUPLC was coupled online through a nanoESI emitter (10 μm tip; New Objective; Woburn, MA, USA) to a Quadrupole-Orbitrap Mass Spectrometer (Q Exactive Plus, Thermo Scientific) using a FlexIon nanospray apparatus (Proxeon). Data was acquired in DDA mode, using a Top10 method. MS1 resolution was set to 70,000 (at 400m/z) and maximum injection time was set to 60 msec. MS2 resolution was set to 17,500 and maximum injection time of 120 msec.

### 2.8 Phosphoproteomics data analysis

Raw data was processed with MaxQuant v1.6.0.16. The data was searched with the Andromeda search engine against the *Fusarium oxyporum* strain 4287 as downloaded from JGI, and appended with common lab protein contaminants. Enzyme specificity was set to trypsin and up to two missed cleavages were allowed. Fixed modification was set to carbamidomethylation of cysteines and variable modifications were set to oxidation of methionines, deamidation of N or Q, and phosphorylation of S, T, or Y. Peptide precursor ions were searched with a maximum mass deviation of 4.5 ppm and fragment ions with a maximum mass deviation of 20 ppm. Peptide, protein and site identifications were filtered at an FDR of 1% using the decoy database strategy (MaxQuant’s “Reward” module). The minimal peptide length was seven amino acids and the minimum Andromeda score for modified peptides was 40. Peptide identifications were propagated across samples using the “match-between-runs” option checked. Searches were performed with the label-free quantification option selected. The quantitative comparisons were calculated using Perseus v1.6.0.7. Decoy hits were filtered out, as well as phosphosites that did not have at least 75% site probability or 2 valid values in at least one experimental group. A Student’s t-test, after logarithmic transformation, was used to identify significant differences across the biological replica. Fold changes were calculated based on the ratio of geometric means of the case versus control samples.

### 2.9 Identifying orthologs

Ortholog identification between genes of *Fusarium oxysporum lycopersici* 4287 and *Saccharomyces cerevisiae* S288C was done by operating a script that performs local BLASTP hit search from protein files of both organisms. The script performs reciprocal best BLAST hit search. When hit is found between query of one organism to the proteome of the other in the first BLAST search, this hit is used in another BLAST search, this time as a query against the reciprocal proteome. As cutoff for the BLAST search parameters of E value of 0.1, length ratio of 0.45 and identity fraction of 0.25 were used.

## 3. Results

### 3.1 DNA repair, protein degradation and transcription modules are up-regulated in response to MMS

To study the cellular response of *F. oxysporum* to continuous exposure of DNA damage, we sequenced RNA from conidia of four independent cultures that were grown for five hours in PDB and then exposed to 0.1% MMS for three hours. We note that, in comparison to previous studies done in *S. cerevisiae*, this is considered a harsh exposure (Jaehnig et al., 2013; Zhou et al., 2016). We also note that, unlike analyses carried out in *S. cerevisiae* and *S. pombe*, the population of conidia is not synchronized to a specific cell cycle (Willis et al., 2016; Zhou et al., 2016). The exposure conditions resulted in survival of 20 % (Fig. S1). We also performed a phosphoproteomic analysis that was done at the same exposure condition and will be described below. RNA sequencing libraries were done using the TruSeq Kit by Illumina and ran by a HiSeq2500 machine. About 95% of reads were uniquely mapped to the genome (for sequencing quality control, see Table S1). We also performed transcriptomic analysis of conidia exposed to half the dose (0.05%) for the same amount of time. This time the libraries made by 3’quantseq method of Lexogen in which only the 3’ of the gene is sequenced (Results for Trueseq methods are shown in Table S2, for 3’quant in Table S3). The mapping rate of 3’ quantseq libraries is less than 85%. Trueseq method identifies differentially expressed genes better than 3’quantseq as determined by side by side analysis of the same RNA (0.1% MMS exposure). Yet, as will be described below the expression of many key genes are differentially induced in lower dose in the same trend as the high dose (Table S3). For the rest of the manuscript we focus on the expression of 0.1% MMS exposure. About 6600 genes were significantly expressed at the RNA level in response to MMS, of which 3563 are up-regulated and 3119 are down-regulated (normalized reads counts and fold changes are found in Table S2).

Classification of the up-regulated genes to gene ontology (GO) groups reveals that DNA repair and protein degradation related genes are significantly enriched (Fig.1 A). Among down-regulated genes, oxidation-reduction and translation-related biological process are significantly enriched (Fig. 1 B). An alternative way to look at the global changes induced by MMS is through identification of cellular networks. To this end, we identified *S. cerevisiae* orthologs of the MMS differentially expressed genes in *F. oxysporum* (see Materials and Methods) and then analyzed these orthologs via STRING network analysis tool (https://string-db.org/cgi/input.pl) (Szklarczyk et al., 2017). STRING identifies protein networks based on evidence for protein-protein interactions. The data input used in our analysis was based on experimental data or databases. Unless stated otherwise the degree of confidence was set to 0.9 (the highest). While the protein networks were built based on data generated in *S. cerevisiae* the evolutionary conservation and significant role of these pathways in the cell suggests that they reflect the biochemical status in *F. oxysporum*. Among up-regulated genes, the modules of proteasome, DNA repair, RNA polymerase II transcription, splicing and processing of ribosomal RNA are highlighted (Fig. 2, Table S4). The picture is similar when analyzing the response to lower MMS dose (Table S3, Fig. S2); specifically the modules of ubiquitin, proteasome and DNA repair are highlighted. Proteasome is highlighted in both concentrations and it is part of nucleotide excision repair (see below) we exposed conidia chronically to MG132 a proteasome inhibitor with and without 0.01% MMS and side by side with exposure to MMS alone. MMS alone does cause delay in growth but as can be seen from Fig. 2 E MMS and MG132 show drug synergy. The combined effect of MMS and MG132 is subtle probably due to pumping out of MG132 as observed in yeast.

**Fig. 1.**
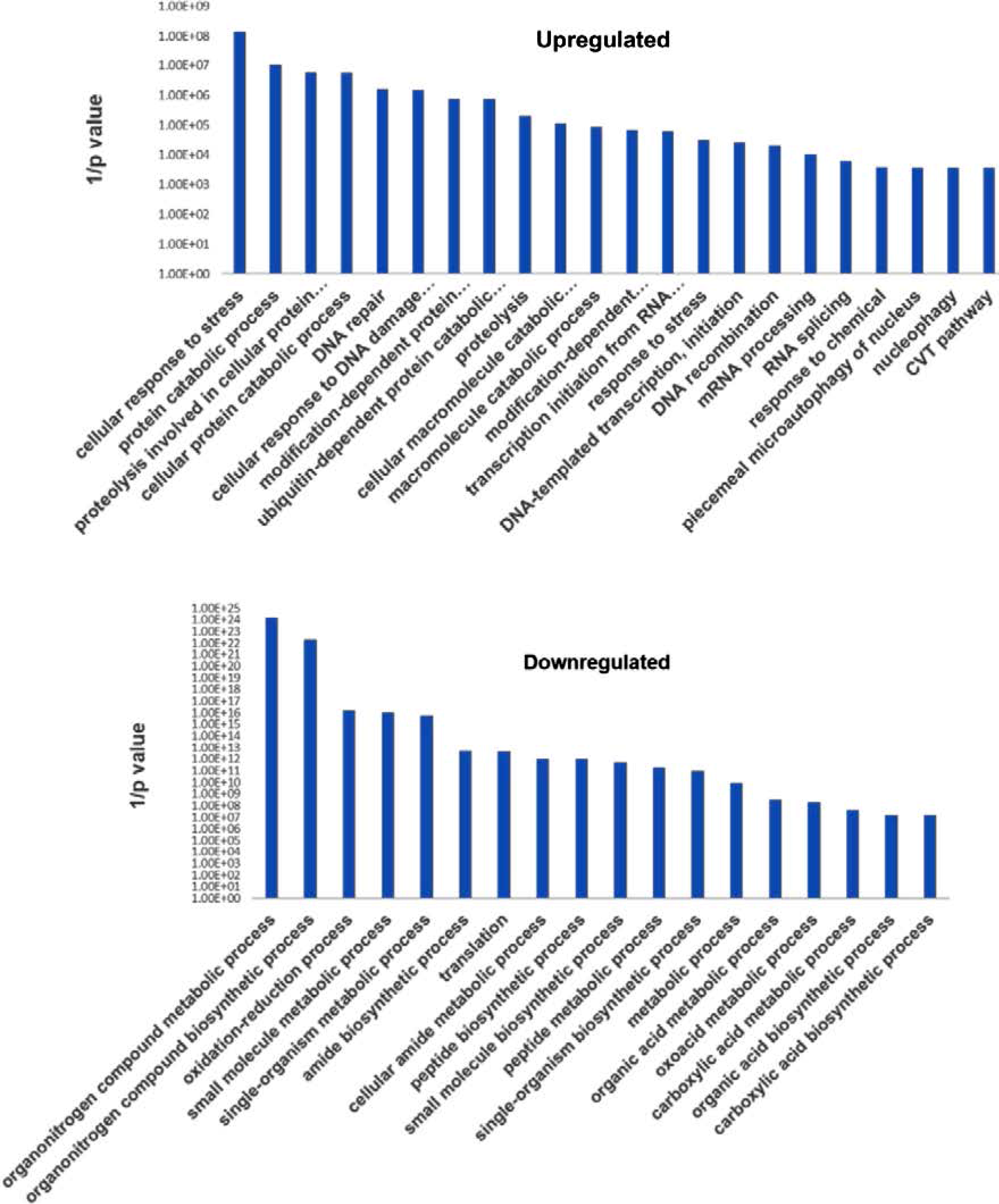
Network analysis of upregulated genes following MMS exposure highlights proteasome, DNA repair and transcription-splicing modules. (A – D). *S. cerevisiae* orthologs of MMS-upregulated *F. oxysporum* genes were identified using reciprocal best BLAST hit analysis (See Materials and Methods). The *S. cerevisiae* orthologs were then analyzed using the STRING platform (https://string-db.org/ (Szklarczyk et al., 2017));. the highest stringency of protein-protein interactions was used (0.9). Experimental data or public databases were used as sources for potential interactions. The number of nodes in the entire network and in each of the sub-networks presented was higher than expected by chance if the entire genome is taken as a reference with a p value lower than 1×10^−16^. Cytoscape was used as a network visualization tool. The most interconnected modules of upregulated genes are presented the model of ribosomal RNA processing is not shown although it is upregulated (for example *sas10*, *sof1* and *utp14, rrb1* see Table S4). E. Conidia were pronged with tenfold serial dilution on plates containing 0.01% MMS, 160, 320 μM of the proteasome inhibitor MG132 (TOCRIS) and 0.01% MMS + 160 μM or + 320 μM of MG132. The plates without MMS were scanned after two nights and the ones with MMS after three nights due to slight growth delay imposed by MMS, this enables to better observe the MG132 effect.

**Fig. 2.**
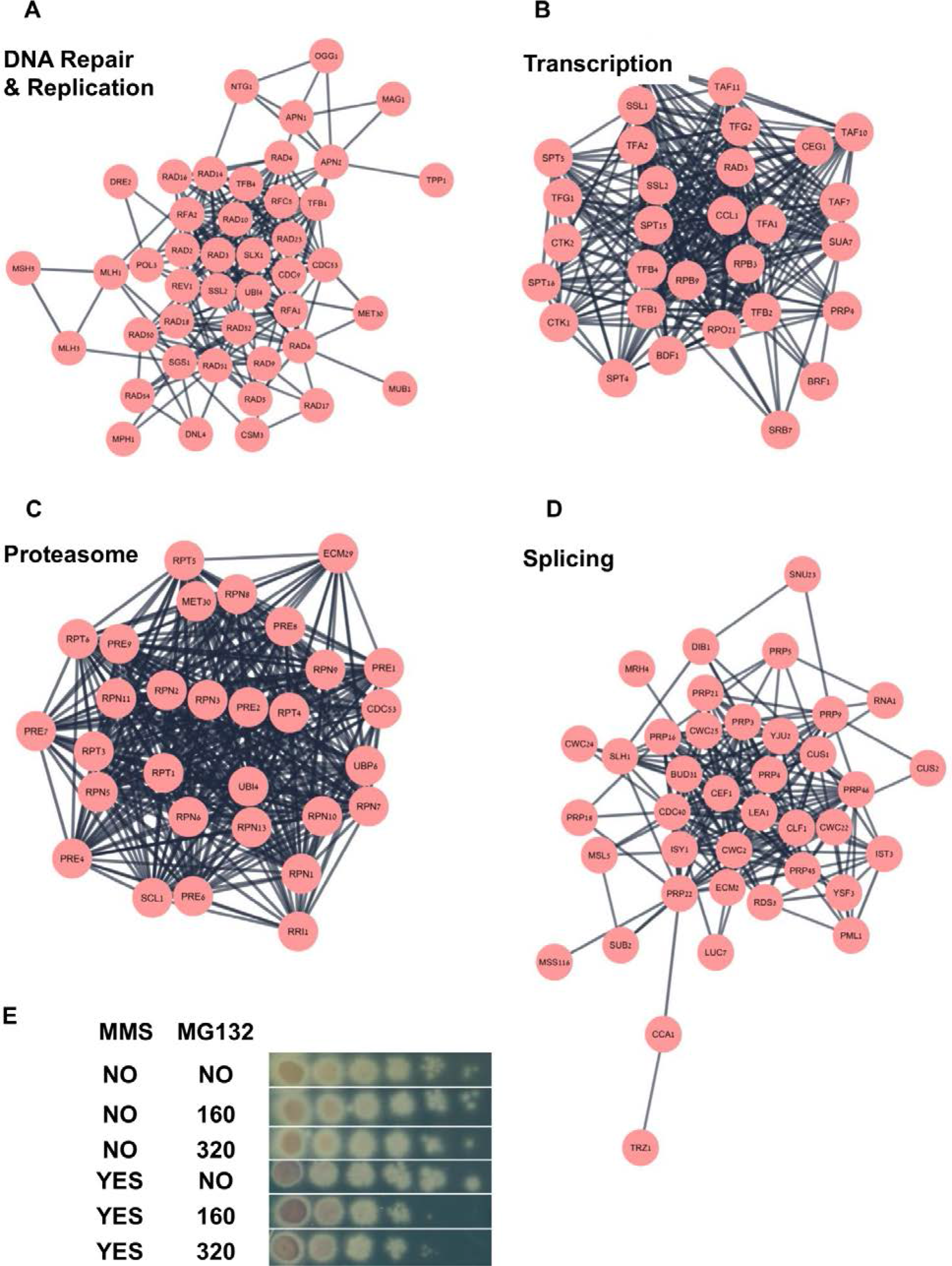
Enriched Gene Ontology terms among differentially expressed genes in *F. oxysporum* following MMS exposure. Classification of up-regulated (A) and down-regulated (B) genes of *F. oxysporum* f.sp *lycopersici* exposed to 0.1% MMS for three hours to gene ontology (GO) terms. GO term finding was done as described under Materials and Methods; differentially expressed genes were identified in Table S3 (2 fold increase < and p < 0.05).

### 3.2 Ribosome biogenesis and respiration modules are down-regulated in response to MMS

Among the down-regulated genes, the ribosomal proteins and mitochondrial/respiration genes are highlighted (Fig. 3, Table S4). In fast growing cells up to 60% of all genes are related to ribosome biogenesis therefore underexpression of ribosomal proteins is a sign of growth arrest (Zhao et al., 2003). Similar picture was observed when the data of conidia exposed to lower dose was analyzed (Fig. S3). We were intrigued by the reduction in mitochondria gene expression; it could be a change in the transcription plan from fast growing to growth arrest. Alternatively, it is possible that mitochondrial DNA is damaged. We hypothesize that chronic exposure to MMS will sensitize cells to respiration inhibitors due to the lower gene expression of respiration-related genes. We exposed conidia chronically to Boscalid, a succinate dehydrogenase inhibitor with and without MMS (SDH2 is down-regulated in our database). We could observe a strong drug synergy between exposure to 0.01% MMS and 40 μg/ml Boscalid (Fig. 3 C).

**Fig. 3.**
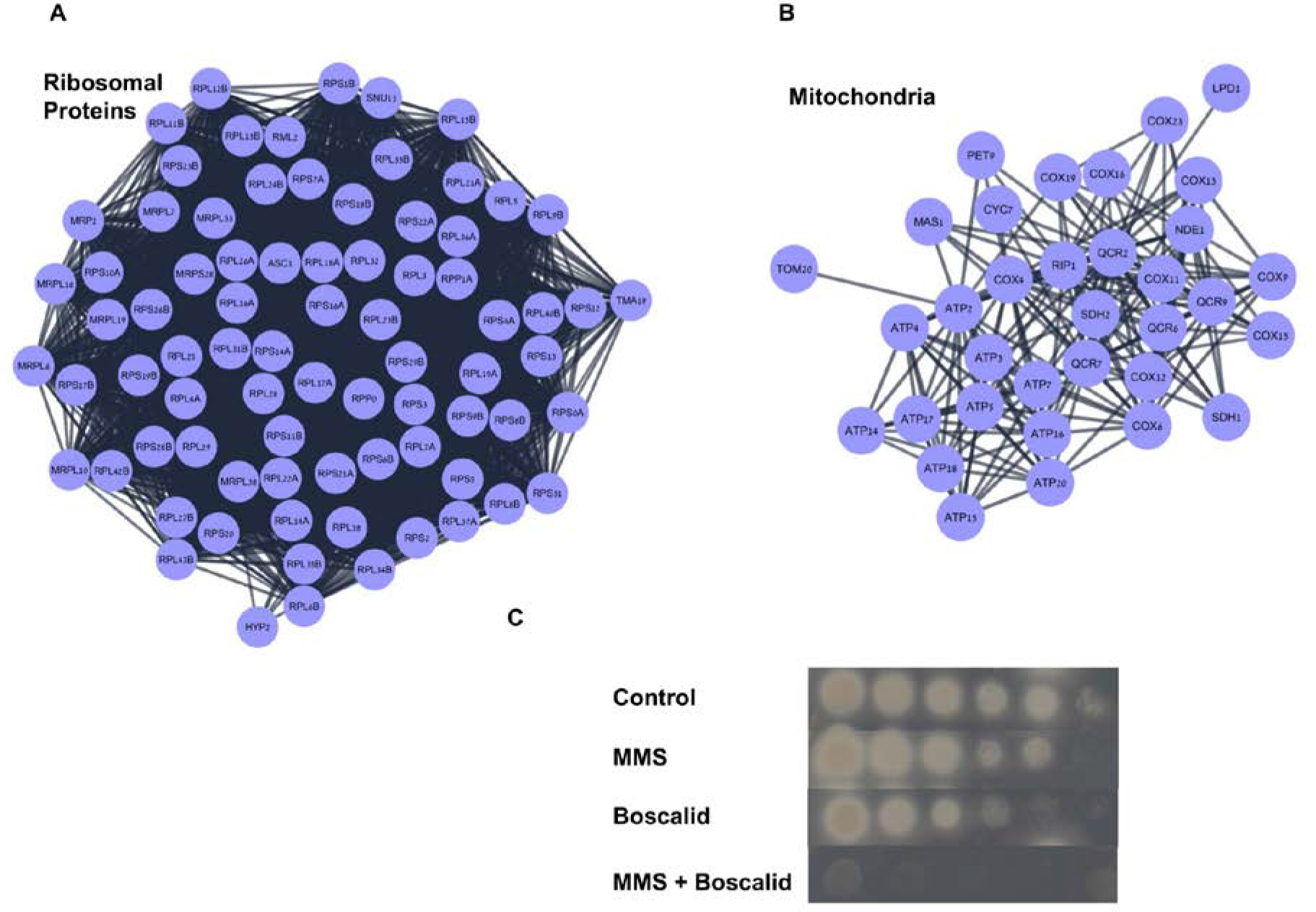
Network analysis of downregulated genes following MMS exposure highlights ribosome, DNA repair and respiration modules. (A, B). *S. cerevisiae* orthologs of MMS-downregulated *F. oxysporum* genes were identified using reciprocal best BLAST hit analysis. The *S. cerevisiae* orthologs were then analyzed using the STRING platform (https://string-db.org/ (Szklarczyk et al., 2017)); the highest stringency of protein-protein interactions was used (0.9). Experimental data or public databases were used as sources for potential interactions. The number of nodes in the database was higher than expected by chance if the entire genome is taken as a reference with a p value lower than 1×10^−16^. Cytoscape was used as a network visualization tool. The most interconnected modules of downregulated genes are presented. C. Conidia were pronged with tenfold serial dilutions on plates containing 0.01% MMS, 40 μg/ml of the succinate dehydrogenase inhibitor Boscalid (Sigma) and 0.01% MMS + 40 μg/ml Boscalid. The plates were scanned after two nights.

### 3.3 *cul4-nedd8-cop9* and global genome nucleotide excision repair are up-regulated in response to MMS in *F. oxysporum*

Upregulation of DNA repair genes in response to MMS is not surprising. The immediate suspects, Mag1 Apn1, Apn2, Parp1 and Mgt1, were strongly up-regulated (Mag1 and especially Mgt1 are highly induced in the lower MMS dose, Table S3). Parp1 is upregulated and phosphorylated in response to MMS (Fig. 4 A) (if fold of increase in phosphorylation is not mentioned, then phosphorylation observed only in treated samples). Base excision repair proteins that are not involved in MMS repair, such as Ogg1 and Ung1, show a mixed response (Fig. 4 A). It’s worthwhile to mention that two Mgt1-like proteins are differentially expressed by MMS; under MMS exposure, one (FOXG_12820) is expressed 800-fold and the second (FOXG_11016) is expressed about 15-fold.

**Fig. 4.**
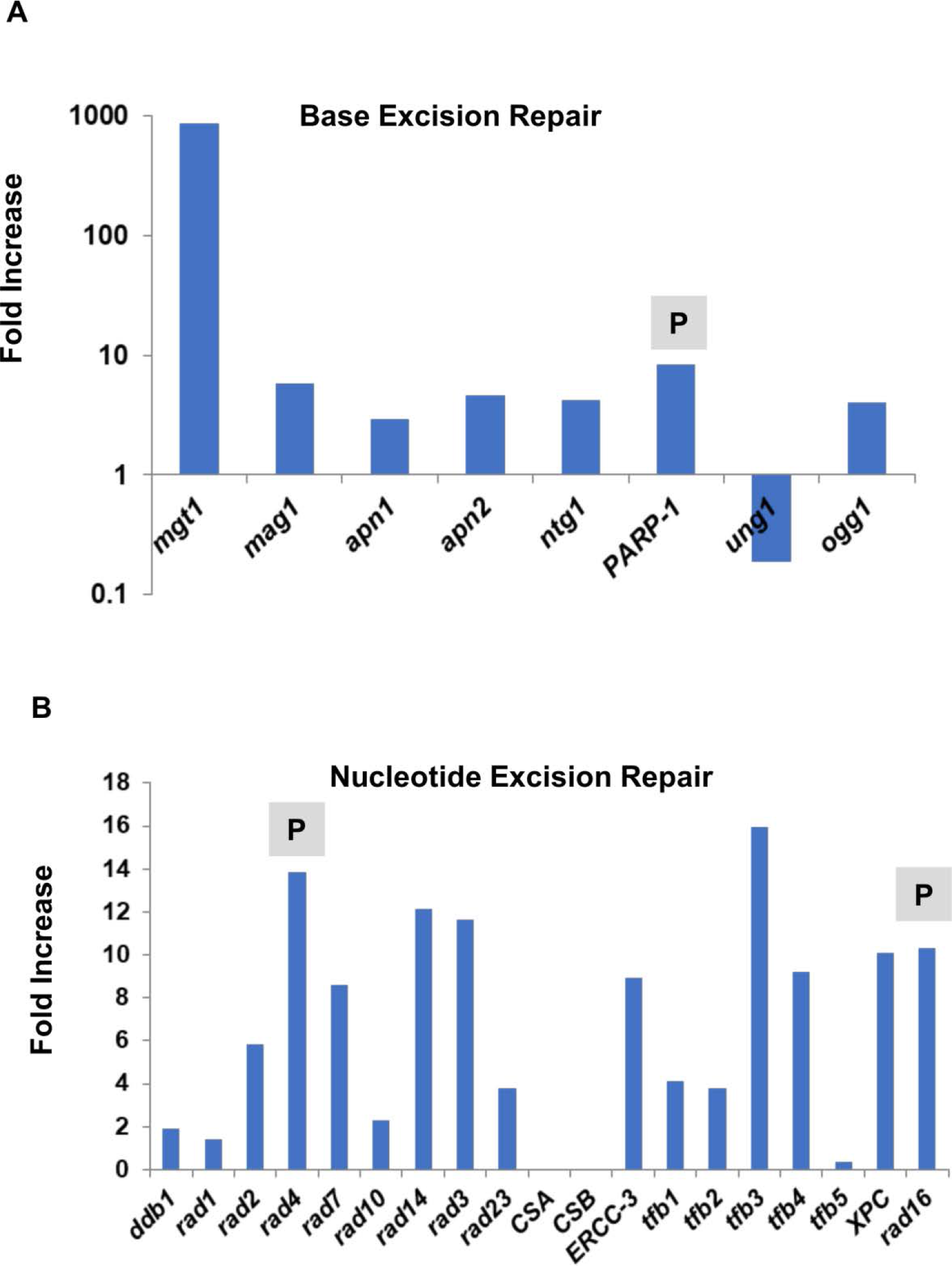
Induction of global genome nucleotide excision repair and homologous recombination both at the RNA and post-translation levels by MMS. The fold increase in the expression of several excision repair genes is presented. Fold increases are calculated from MMS and control-adjusted RNA sequencing reads with *p* < 0.05. (A) Base excision repair genes; (B) Nucleotide excision repair genes. P - indicates for phosphorylated proteins following MMS exposure (> two-fold, *p* < 0.05).

Somewhat surprising was the concerted up-regulation of nucleotide excision repair genes. Interestingly, this is mainly due to activation of the global genome repair (similar results were shown for low dose exposure, Table S3). Rad16 and Rad2, two nucleases of NER, are also phosphorylated (Fig. 4 B). Importantly, we could not measure the expression of transcription coupled repair (Fig. 4 B). As mentioned above, we found that modules of protein ubiquitination and modifications, as well as proteasomes, to be over represented among up-regulated genes (Fig. 1 Fig. 2, Fig. S2). A closer look revealed that genes that function in protein degradation and NER are up-regulated in response to MMS. That includes Ubiquitin activating enzyme (E1) (Uba1, FOXG_08381); Ubiquitin conjugating enzyme (E2) (Ubc12, FOXG_05289); Ubiquitin ligases (E3) (RBX1/HRT1, FOXG_10518) and E3 co-factor (Cul4, FOXG_09372). These proteins are part of the of theCop9- Nedd8 pathway that is degrades the NER proteins Ddb1 and Ddb2 (Fig. S4) (Chung and Dellaire, 2015). In addition, we observe activation of Cul3 that is needed for degradation of stalled RNA polymerase II (see Discussion section). Several Cul4, Cul3 and proteasome related genes are up-regulated also in the response to lower MMS dose (Table S3). Apart from NER, the proteasome system is up-regulated altogether including ubiquitin itself, three ubiquitin E1 enzymes, 11 E2s, and 18 E3s ubiquitin/sumo enzymes. In addition, 30 proteasome subunits are upregulated. Eight enzymes that recycle or break the bond of ubiquitin with its target are also upregulated. In the context of DNA damage, the ubiquitin E3 Rad18 and the sumo ligase Wss1 are activated by MMS either at the transcription or the phosphorylation level (Table S5).

The homologous recombination pathway is up-regulated in a concerted manner, as can be seen in Fig. 5. Genes involved in recombination initiation (Mre11, Rad50); joint molecule formation (Rad52, Rad54, Rad51, Rad55) and recombination intermediates resolution Mus81 and Sgs1) are up-regulated. Rad52 and Sgs1 are also phosphorylated in response to MMS exposure. Several recombination related genes are shown to be up-regulated in response to lower MMS exposure (Table S3, Fig. S2). In addition, the non-homologous end joining (NHEJ) machinery is also up-regulated in response to MMS (Fig. 5). Homologous recombination repairs breaks in the DNA and also helps to rescue stalled replication forks. Post-replication repair is also a major player in this process. Post-replication repair consists of two branches: error free, through template switching, and error-prone, mediated by translesion DNA polymerases. *rad5* is the major template switching gene is up-regulated by MMS (Fig. 5). It is not clear which of the translesion DNA polymerases contributes to creating mutations in response to MMS exposure in *F. oxysporum*. Nevertheless, it is interesting that the expression of *rev3*, the polymerase responsible for most of the MMS induced mutations in fungi, is actually reduced (Fig. 5) (Sakai et al., 2002; Conde and San-Segundo, 2008). The picture is complex for other translesion DNA polymerases: the expression of *rev1*, polymerase kappa and polymerase iota is increased, while *rad30* is decreased. We next studied the mutagenic effect of the exposure by determining the frequency of benomyl resistant isolates with and without exposure (same MMS exposure as done for RNA sequencing). Interestingly, we observed only modest induction in mutation frequency; 6×10^−5^±3×10^−5^ for treated samples vs 2×10^−5^±0.5×10^−5^ for untreated one. This 3 fold induction of mutation frequency is even lower than the effect on survival (Fig. S1).

**Fig. 5.**
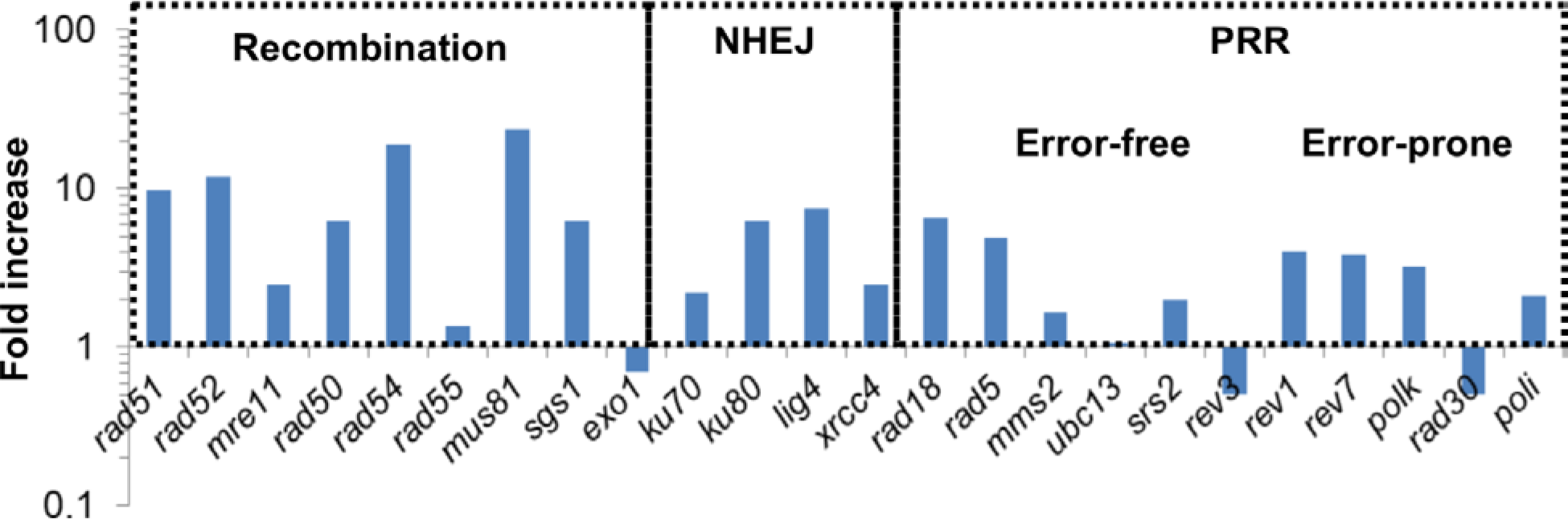
Induction of homologous recombination and post-replication repair genes both at the RNA and post-translation levels by MMS. The fold increases in the expression of several double strand break repair and post-replication repair genes are presented. Fold increases are calculated from MMS and control-adjusted RNA sequencing reads with *p* < 0.05. P-indicates for phosphorylated proteins following MMS exposure (> two-fold, *p* < 0.05).

To sum up activation of DNA repair pathway by MMS in *F. oxysporum*, activation of DNA repair genes occurs mainly at the transcription rather than phosphorylation level. Other protein modifications, such as neddylation, ubiquitination and sumoylation may play an important role here. While expression of several error-free repair mechanisms are significantly induced by MMS, error-prone related genes are mildly induced or even reduced.

### 3.4 MMS activates the Yap1, Xbp1 transcription modules

We observe a massive transcriptional response to MMS that strongly regulates key genes in DNA repair (Fig.s 1, 2, 3, 4). Therefore, we identified transcription factors that are either up-regulated or phosphorylated in response to MMS exposure. About 150 transcription factors are up-regulated by MMS and 122 are phosphorylated (See Table S6).

Yap1 was found to be up-regulated at the transcription level (four-fold, it is also up-regulated at lower doses Table S3); it is also phosphorylated in response to MMS exposure. Yap1 is an immediate candidate to be a master regulator of the DNA damage response in *F. oxysporum*. Based on previous experimental results summarized at the *Saccharomyces* Genome Database, we identified the *F. oxysporum* orthologs of Yap1 target genes (Cohen et al., 2002; MacIsaac et al., 2006; Hu et al., 2007; Venters et al., 2011); more than half of the orthologs found (153/287 higher induction comparing to the entire genome *p*-value is < .00001) were differentially expressed by MMS; 60 were up-regulated and 93 were down-regulated. Among up-regulated genes, we found important players of the major transcriptional module that are activated by MMS, including recombination proteins Rad51 and Rad52, nucleotide excision repair Rad4, ubiquitin, ubiquitin conjugating and proteasome subunits (Ubi4, Rad6 and Rpn11, respectively) (Fig. 6, Table S7). Among the down regulated genes, 23 were related to ribosomal proteins; it seems that Yap1 is a major regulator of these proteins (Fig. 6, Table S7).

**Fig. 6.**
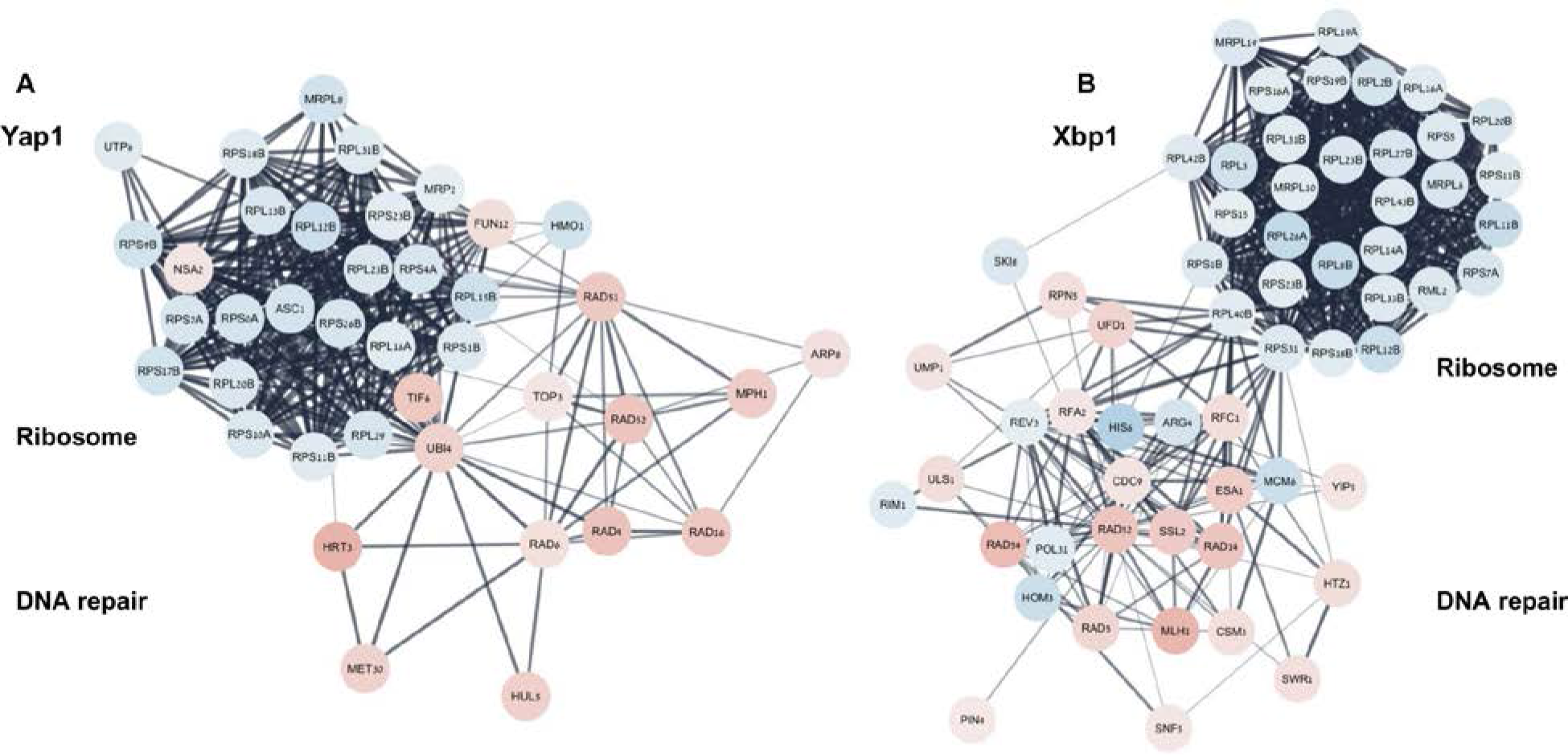
*Fusarium oxysporum* Yap1 and Xbp1 transcriptional networks are activated by MMS. *F. oxysporum* orthologs of *S. cerevisiae* genes that are under Yap1 regulation as previously determined by expression and chromatin IP experiments in *S. cerevisiae* were identified. Genes that are either induced or suppressed following exposure of *F. oxysporum* to MMS were further selected to be analyzed by the STRING platform (as described in the legend to Fig. 2). Networks of up-regulated genes are marked in red and down-regulated marked in blue. The color gradient represents the fold increase (brightest is the strongest effect). (B) Same as (A), but for Xbp1. Mgt1 and Ddi3 that are significantly induced by MMS and are known to be under Xbp1 control in *S. cerevisiae* are not part of the network. The number of nodes in both networks is higher than expected by chance (p value < 1×10^−16^).

Xbp1 is a protein that in *S. cerevisiae* is not expressed in log phase cells, but is expressed under stress conditions (Miles et al., 2013). We performed for Xbp1 the same analysis done for Yap1 based on previous experimental data obtained in *S. cerevisiae* (MacIsaac et al., 2006; Venters et al., 2011); more than half of the orthologs found (343/ 668 higher induction comparing to the entire genome *p*-value is < .00001) were differentially expressed by MMS; 152 were up-regulated and 191 were down-regulated. Among up-regulated genes were the most *F. oxysporum* responsive genes to MMS, Mgt1 and Ddi3. Some of the over and down-regulated genes are shared with Yap1 in agreement with the fact that in *S. cerevisiae*, Xbp1 is found upstream of Yap1 (Fig. 6 Table S7).

We found many transcription factors of which a clear ortholog was not found in *S. cerevisia*e. An example of a very interesting transcription factor that is MMS-induced over 100-fold is FOXG_17458. It is encoded on the accessory/pathogenicity chromosome 14, and when up-regulated, activates several virulence genes (*SIX* 1-14) (Table S6) (van der Does et al., 2016). However, in our database the *SIX* genes are not induced indicating that natural up-regulation alone of this gene is insufficient to drive the expression of pathogenicity genes. Chromosome 14 is known as lineage specific (LS), a chromosome that is not found in the genomes of taxa closely related to *F. oxysporum lycopersici*. There are other regions in the *F. oxysporum* genome that are LS those regions are often associated with pathogenicity. We examined the effect of MMS on genes encoded on LS regions and found out 143 genes that are up-regulated and seven genes that are phosphorylated (Table S6).

### 3.5 DNA damage checkpoint activation in *F. oxysporum* is mainly manifested by Chk1 and Chk2 phosphorylation

We also study the changes in phosphoproteome following exposure of conidia from three cultures to 0.1% MMS for three hours. Samples for phosphoproteomic analysis were prepared as described under Materials and Methods; details of the proteomic technical specifications are found in Table S8. 823 proteins were phosphorylated differentially and 302 proteins were de-phosphorylated differentially in response to MMS (Table S8, cutoff phosphorylation status was two fold increase or decrease from the untreated reference with p value lower than 0.05). Enriched GO terms of biological processes of phosphorylated proteins were cellular localization, cellular processes, cellular component biogenesis, gene expression and intracellular transport (Fig. S5 A). Enriched GO terms of biological processes of de-phosphorylated proteins did not indicate for a clear pathway (Fig. S5 B). The change in gene expression and phosphorylation status for DNA damage checkpoint proteins was examined. Unlike DNA repair, activation on the transcription level was modest (up to four-fold). The ortholog of the *S. cerevisiae RAD9* gene and two of orthologs of the *S. pombe* 911 complex are up-regulated in response to MMS. The same is true for the DNA replication stress checkpoint Csm3 (Table 1). MMS-dependent phosphorylation was only observed for the two DNA damage transducer kinases Chk1 and Chk2. Interestingly, phosphorylation of ATM or ATR was not observed (Table 1). This could stem from technical reasons or it could be that autophosphorylation of these proteins does not occur in *F. oxysporum*. The GTPase Bub2, part of the mitosis network, is both phosphorylated on S43 and dephosphorylated on S91 in response to MMS. As described below several cell cycle regulators are phosphorylated in response to MMS.

**Table 1.** Effect of MMS on DNA damage signal transduction and checkpoint proteins in *F. oxysporum*. The effect of MMS on gene expression and phosphorylation of proteins that are part of DNA damage signal transduction and DNA damage checkpoint pathways was determined. The gene description was taken from the Saccharomyces Genome Database. FOL ID is of *F. oxysporum lycopersici* 4287. Fold increase RNA is the ratio of treated to untreated. Phosphorylation values represent the ratio between treated to untreated. Unique to MMS means that phosphorylation was observed only in treated samples and unique to controls means that phosphorylation was observed only in untreated samples (see Table S8). Complex phosphorylation status means some residues are phosphorylated following the treatment and others are de-phosphorylated. ND- not determined by the mass-spec analysis.

| Gene | Description | FOL ID | RNA fold increase | Phosphorylation |
| --- | --- | --- | --- | --- |
| *rad9* | DNA damage signal transduction adaptor | FOXG_07943 | 3 | ND |
| *rad24* | 911 checkpoint complex | FOXG_00420 | Not enough reads | No significant change |
| *rad17* | 911 checkpoint complex | FOXG_01178 | 3.3 | ND |
| *mec3* | 911 checkpoint complex | FOXG_07970 | 2.8 | ND |
| *mrc1* | S phase checkpoint protein | FOXG_05485 | 0.4 | Unique to MMS |
| *atr* | Protein kinase, DNA damage signal transduction | FOXG_18483 | 1.8 | ND |
| *chk1* | Protein kinase, DNA damage signal transduction | FOXG_11222 | 1.6 | Unique to MMS |
| *chk2*<br>(*rad53*) | Protein kinase, DNA damage signal transduction | FOXG_01680 | 1.6 | Unique to MMS |
| *atm* | Protein kinase, DNA damage signal transduction | FOXG_20040 | 0.6 | ND |
| *bub1* | Protein kinase, anaphase checkpoint | FOXG_00034 | 1.9 | ND |
| *bub2* | Mitotic exit network regulator | FOXG_00324 | 1 | Complex |
| *bub3* | Kinetochore checkpoint protein | FOXG_08492 | 1 | ND |
| *mad1* | Spindle assembly checkpoint | FOXG_09116 | 0.6 | 2.85 |
| *mad2* | Spindle assembly checkpoint | FOXG_10615 | 0.2 | ND |
| *csm3* | DNA replication checkpoint | FOXG_15624 | 3.4 | ND |
| *bim1* | Mitosis checkpoint | FOXG_02318 | 1 | No significant change |
| *elg1* | DNA replication checkpoint | FOXG_10311 | 0.3 | ND |

### 3.6 Three different MAPK pathways and cAMP signaling are modulated in response to MMS

Until now, we focused on DNA damage related genes, their transcriptional regulators and activators. However, our analysis revealed activation of other stress/signaling pathways. We examined the *F. oxysporum* kinase repertoire for kinases that are phosphorylated in response to MMS (DeIulio et al., 2018). 34 kinases are phosphorylated by MMS (Table S9). Using the KEGG pathway database (http://www.genome.jp/kegg/pathway) (Kanehisa et al., 2017) we were able to identify three MAPK pathways modulated by MMS. Components of the MAPK pathway involved in cell wall integrity are phosphorylated from the cell membrane (Wcs1) to the nucleus (Rlm1) (Fig. 7, Table S9). The pheromone/starvation response is also modulated by MMS. The membrane-bound Yck1 is phosphorylated, as well as Ste20 and Ste11 MAPK. Our data indicate that the osmolarity stress is influenced by MMS. This is because Ssk2, Pbs2 and Hog1 are phosphorylated (Fig. 7, Table S9).

**Fig. 7.**
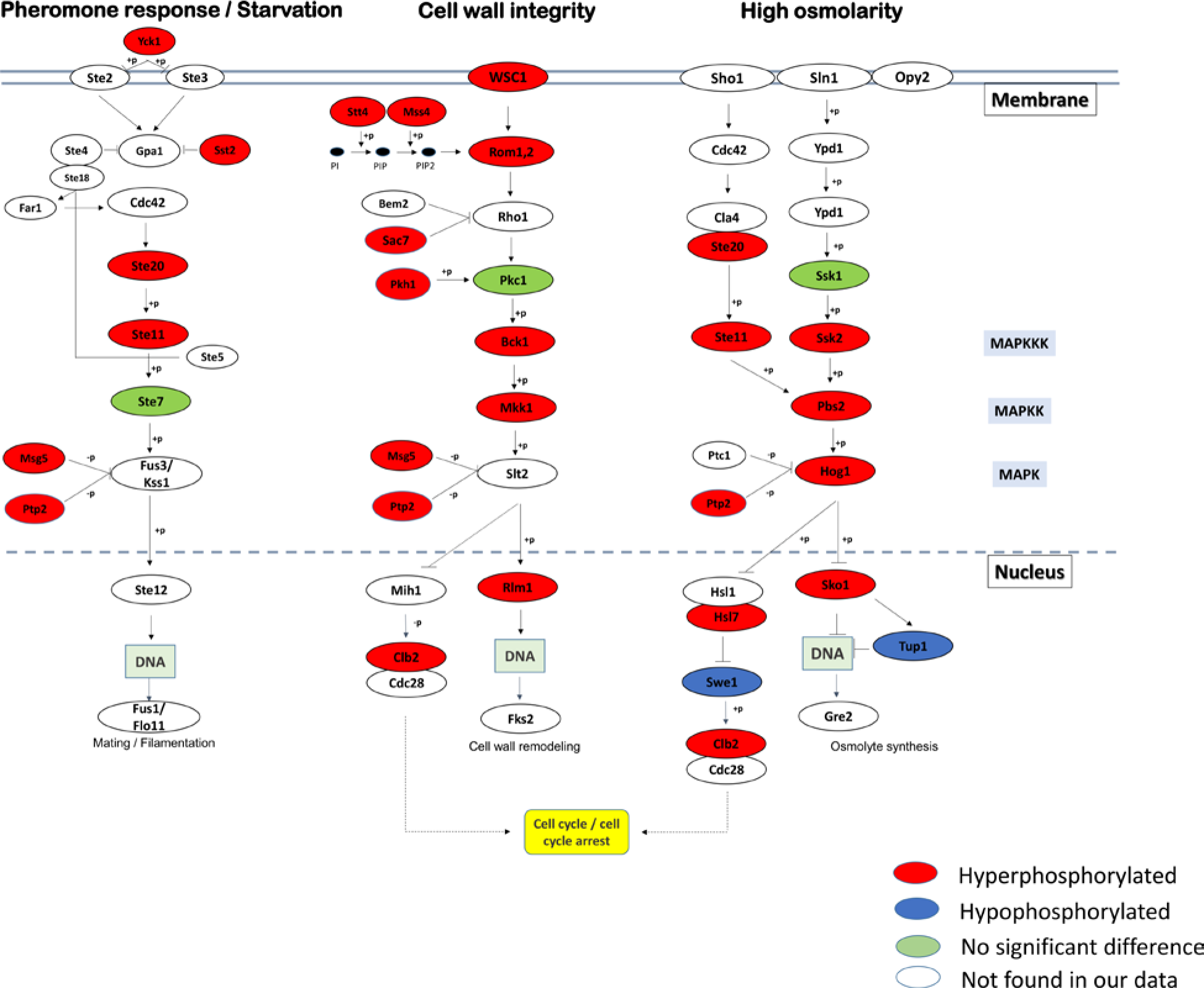
Activation of three MAPK pathways by MMS in *F. oxysporum*. MAPKs and components of MAPK signaling pathways were identified among phosphorylated proteins following MMS exposure (> two-fold, *p* < 0.05). MAPKs were identified using KEGG and a list of kinases of *F. oxysporum* (Li Jun Ma personal communication).

Using STRING analysis we were able to identify several proteins that serve as hubs in the phosphorylated database (analysis was done as described above and in the legend to Fig. 2). The most interesting hub was centered on Cyr1, adenylate cyclase, the enzyme that generates cAMP in the cell (Fig. 8 A, Table S10). Associated with Cyr1 was Sgt1, a Hsp90 co-chaperone that is involved in kinetochore formation and cell cycle progression (Kitagawa et al., 1999; Catlett and Kaplan, 2006). Other cell cycle regulators that connect with Cyr1 and phosphorylated by MMS are the splicing factor Cdc40 and Dma2. Other cell cycle regulators are also phosphorylated by MMS, for example Cdc14 is found within a network of chromosomal proteins such as Rad52, and topoisomerase 2 (Fig. 8 B, Table S10). Cdc4 and Bub2 are cell cycle regulators that are phosphorylated but high confidence STRING analysis did not place them in the Cyr1 network. According to the STRING analysis, Cyr1 is interconnected to several signal transduction pathways that are activated by MMS in our phosphoproteomic dataset. The Snf1 pathway (both Snf1 and Snf2 are phosphorylated by MMS in *F. oxysporum*); several MAPK pathways are connected to Cyr1. In the Hog1 pathway, Pbs2 and Hog1 interact with Cyr1 and are activated by MMS, as seen in Fig.s 6 and 7 A. In the starvation/pheromone signaling pathway, Ste11 interact with Cyr1, as seen in Fig.s 6 and 7 A. Another module of phosphorylated proteins is rRNA processing that is also upregulated at the expression level, (Fig. 2, Fig. 8 C, Table S10).

**Fig. 8.**
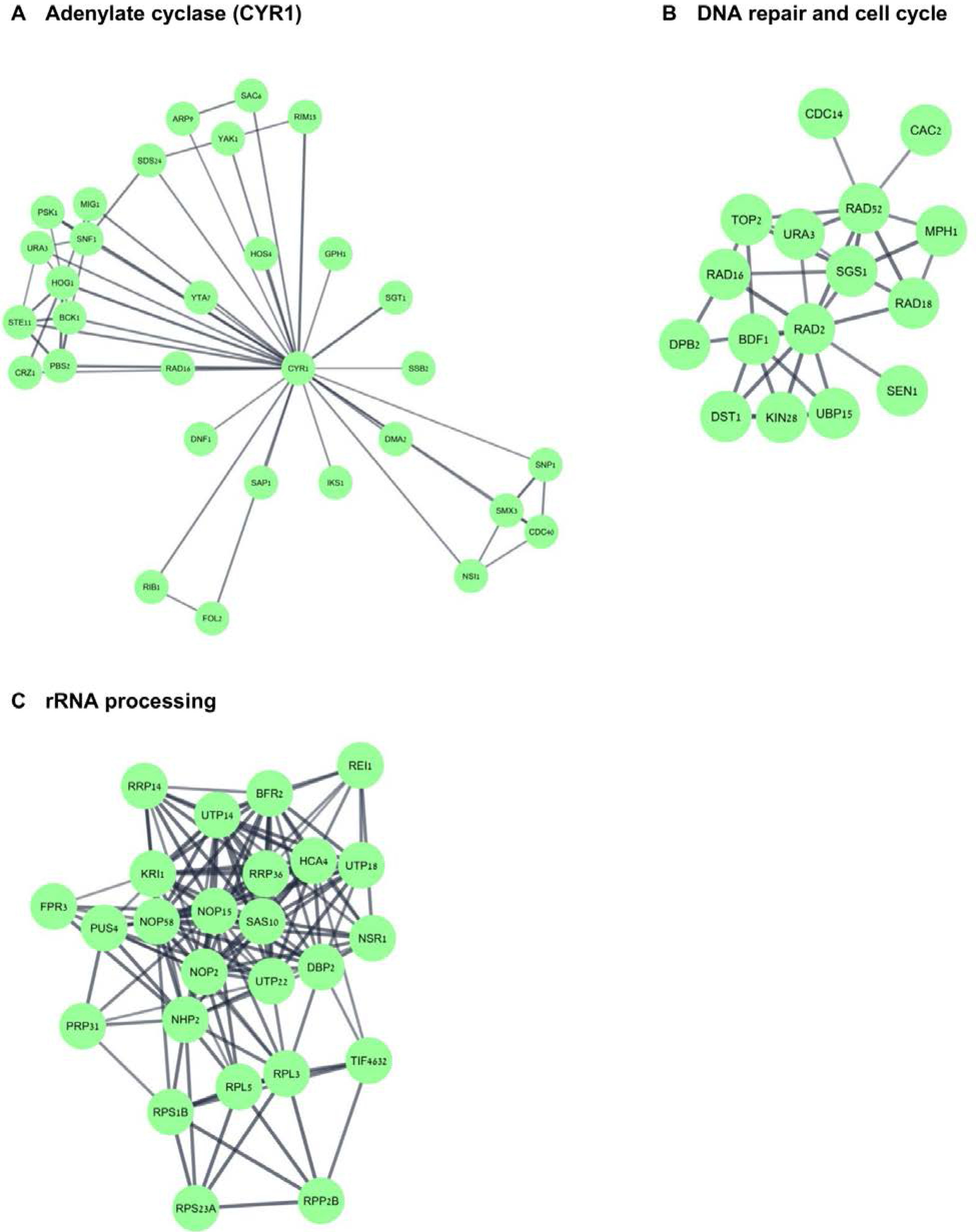
Adenylate cyclase forms a hub of MMS-induced phosphorylated proteins. *S. cerevisiae* orthologs of *F. oxysporum* MMS-induced phosphorylated proteins (> two-fold, p < 0.05) were analyzed using STRING to identify cellular modules with a 0.7 confidence score. (A). Cyr1 network. (B) Cell cycle and DNA repair network. (C) rRNA processing network. The number of nodes in both networks is higher than expected by chance (p value < 1×10^−16^).

## 4. Discussion

### 4.1 Comparison with *S. cerevisiae* at the system level

We discuss our results from a comparative genomic stand point. We compared our results to several publications focused on *S. cerevisiae* and *S. pombe*. In *S. pombe*, gene expression analysis of MMS treated cells was done in far milder conditions and did not reveal a strong DNA repair response (Chen et al., 2003; Lackner et al., 2012). Shalem et al. performed an RNA expression analysis of *S. cerevisiae* cells in similar conditions to here (0.1% MMS for three hours (Shalem et al., 2008)). We compared the list of *S. cerevisiae* orthologs that are up-regulated in *F. oxysporum* to the one published by Shalem et al., (Shalem et al., 2008). Out of 1005 up-regulated genes in *S. cerevisiae* 831 genes are found to be *S. cerevisiae* specific namely no *F. oxysporum* orthologs were over expressed by MMS STRING analysis of these genes included several relatively small modules that contained for examples members of the cyclosome and histone modification enzymes (data not shown). 174 genes were shared between *S. cerevisiae* and *Fusarium*; STRING analysis of these genes revealed that the modules of the proteasome and several DNA repair genes are highlighted (Fig. 9). 545 genes were up-regulated in *F. oxysporum* but not in *S. cerevisiae*. These genes can be divided to two groups. First, genes that belong to a module that is up-regulated in *S. cerevisiae* such as the proteasome or DNA repair (i.e more of the same). Second, genes that form modules that are not MMS responsive in *S. cerevisiae* including transcription, splicing and ribosomal RNA processing (Fig. 9). DNA damage interferes with transcription, to overcome this obstacle cell operates transcription coupled repair; we do not observe activation of transcription coupled repair here (Fig. 4). A last resort solution to damage-stalled RNA polymerase is degradation of RNA pol II and transcription restart (Wilson et al., 2013). Only in *F. oxysporum* did we observe up-regulation of RNA polymerase II degradation machinery (the Cul1,2,3 system) and RNA pol II associated proteins (*tfa1*, *taf11*, *spt5*) (Table S4, Table S11 and Fig. 9). Since in *F. oxysporum* introns are more common than *S. cerevisiae* it is reasonable that induction of the transcription machinery will be followed also by induction of the splicing machinery (Fig. 9). We still do not understand why genes involved in ribosomal RNA processing such as *sas10*, *sof1* and *utp14* are up-regulated following MMS exposure while ribosomal proteins are down regulated (Fig. 2Fig. 3). DNA damage might interrupt with ribosomal RNA transcription; we suggest that to coordinate proper ribosomal assembly *F. oxysporum* cells have to both up regulate the RNA processing and down regulate the amounts of ribosomal proteins. The module of mitochondria and respiration is down-regulated in *F. oxysporum* but not in *S. cerevisiae* following MMS exposure (Table S11, Fig. 3). *S. cerevisiae* rely more on oxidative glycolysis to generate energy than other fungi and therefore probably growth arrest does not affect the expression of respiration related genes so much (Hagman et al., 2013). Interestingly, chronic exposure to MMS strongly synergizes with Boscalid a succinate dehydrogenase inhibitor; in agreement with reduction in the expression of *sdh2* following MMS acute exposure. We hypothesize that change in the transcription plan following MMS exposure is enough to sensitize *Fusarium oxysporum* to Boscalid.

**Fig. 9.**
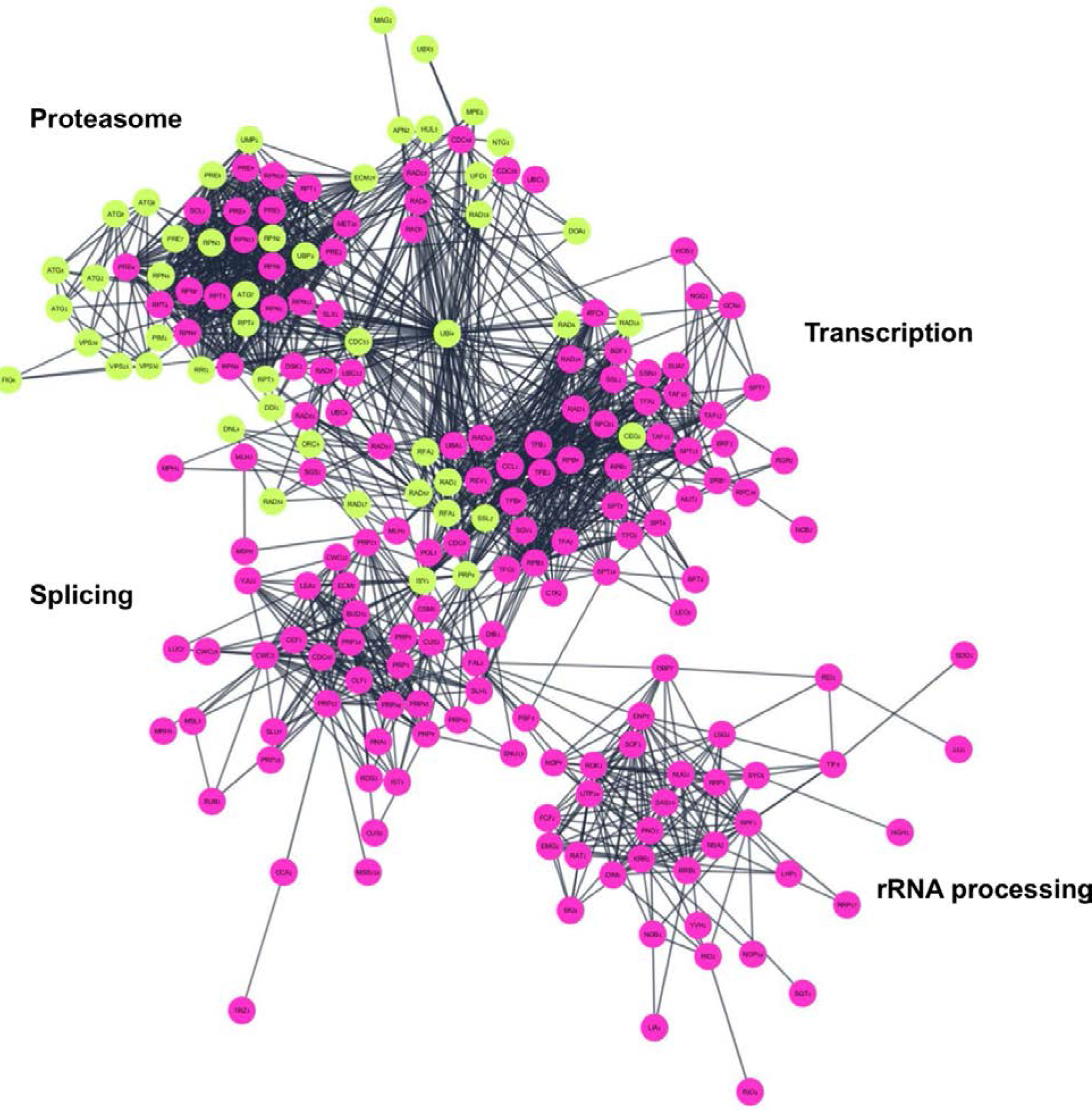
Comparison between S. cerevisiae and *F. oxysporum* up-regulated genes from a system standpoint. Overlap between MMS-upregulated *S. cerevisiae* genes (Shalem et al., 2008) and *S. cerevisiae* orthologs of MMS-upregulated *F. oxysporum* genes. STRING analysis was conducted as in Fig. 2. The number of nodes in both networks is higher than expected by chance (p value < 1×10^−16^). In pink proteins that are up-regulated both in *S. cerevisiae* and *F. oxysporum* in blue proteins that are expressed only in *F. oxysporum*. The proteins in each module are identical to the ones presented in Fig. 2.

A major pathway involved in the response of *F. oxysporum* to MMS is the Cop9-Nedd8-Cul4 pathway (Table S5, Fig. S4). This pathway leads to protein degradation; in yeast the pathway is incomplete several proteins including Cul4 are not encoded. Protein degradation was identified long ago to be involved in the response to MMS, probably because MMS also affects proteins (Fry et al., 2005). In addition, RNA polymerase II that is stalled by DNA damage is ubiquitinated and degraded as described above (Wilson et al., 2013). Our hypothesis is that activation of the Cop9-Nedd8-Cul4 pathway is the reason why there is overactivation of ubiquitin/proteasome related genes in *F. oxysporum* vs *S. cerevisiae*.

### 4.2 DNA repair and DNA damage signal transduction

We observed that the global branch of the nucleotide excision repair (NER) pathway was activated by MMS (Fig.s 1, 3 S4). There have been previous documentations of activation of the NER system by MMS at the transcription level, but to a lesser extent (Benton et al., 2006; Shalem et al., 2008). The major difference between the *S. cerevisiae* and the *F. oxysporum* activation of NER is the involvement of the Cop9- Nedd8-Cul4 pathway, which does not exists to the same extent in *S. cerevisiae* (Fig. 5) (Li et al., 2011). This difference highlights the fact that activation of NER as a system is evolutionary conserved in MMS damage. While NER is considered to be a repair pathway of bulky DNA lesions, previous *in vitro* and *in vivo* experiments had shown it also may be involve in repair of MMS lesions (Huang et al., 1994; Memisoglu and Samson, 2000; Sancar and Reardon, 2004). While we see a clear induction at the mRNA level of DNA repair proteins, MMS-induced phosphorylation was not as significant (Table 1, Fig.s 3 4). We do not see phosphorylation of mismatch repair proteins, nuclear pore proteins and Mre11, Rad9 and Mrc1 as recently reported in *S. cerevisiae* (Zhou et al., 2016). In general, several DNA damage repair proteins that are phosphorylated in *S. cerevisiae* following MMS exposure are up-regulated in *F. oxysporum* (Table 1, Fig. 2, 4, 5). Like *S. cerevisiae*, we observe phosphorylation of Rad52, Sgs1, Parp1, Chk1, Chk2, topoisomerase I and II (Table 1, Fig. 4, 5, 8).

The expression comparison revealed an interesting difference regarding strategy of repair. In *F. oxysporum* the *mgt1* gene was induced by 800-fold and another paralog 15-fold, 0.01% MMS exposure caused induction of 32 fold in Mgt1(Fig. 4, Table S3). The induction of the *S. cerevisiae* ortholog under the same MMS exposure was far less dramatic although a slight induction in the expression (up to two-fold) was observed (Benton et al., 2006; Shalem et al., 2008; Jaehnig et al., 2013). In contrast, *mag1* was up-regulated to a similar level or even higher in *S. cerevisiae* (three to 30-fold increase in *S. cerevisiae* versus five-fold in *F. oxysporum*).

### 4.3 Stress signaling

The MMS exposure applied here generated significant stress; genotoxic and proteotoxic stresses are expected as well as indirect effect due to growth arrest. The MAPK signaling of cell wall integrity, osmolarity stress and pheromone/starvation stresses are activated (Fig. 7). Activation of MAPK in response to DNA damage and the role of its activity in DNA damage response was shown in other organisms (Watson et al., 2004; Willis et al., 2016) (and the references therein). However, we could not observe the same level of MAPK pathway activation through phosphorylation in neither the *S. pombe*, nor in the *S. cerevisiae* phosphoprotein dataset (Willis et al., 2016; Zhou et al., 2016). This could stem from the differences in the experimental set-up or technical aspects of the specific proteins involved. Thirty four protein kinases were among proteins called as significantly differentially phosphorylated. We saw enrichment in the AGC, CAMK, and CK1 and depletion of the Histidine and Atypical kinase families among differentially phosphorylated kinases, consistent with the fundamental roles of the AGC, CAMK, and CK1 kinases in the regulation of cell cycle and growth and development. The lineage specific genome of Fo4287 has been shown to contribute to the expansion of kinase families beyond the core genome and contributes to the differences among FOSC strains (described in (DeIulio et al., 2018))). We identified seven genes within the phosphoproteomic data which were predicted to belong to lineage specific contigs, none of which were kinases. Signaling and responding to DNA damage is a fundamental process common to all organisms and the exclusive role of the Fo4287 core genome protein kinases in this response is consistent with a core genome which controls basic life functions within *F. oxysporum*.

We observed phosphorylation of adenylate cyclase following MMS exposure. Moreover, adenylate cyclase was shown to be connected to many other proteins that are phosphorylated in response to MMS (Fig. 8 A). Using the same tools as used for *F. oxysporum* we analyzed the data described by Zhou et al., and found that adenylate cyclase forms a hub of MMS phosphorylated proteins also in *S. cerevisiae* (Zhou et al., 2016). Previously a connection between cAMP signaling and DNA damage dependent cell cycle arrest was made in *S. cerevisiae* (Searle et al., 2004). Adenylate cyclase affects many pathways in the cell. In filamentous fungi it affects normal growth, sporulation, and plant disease; it is also an anti-stress-response protein (Ivey et al., 2002; Choi and Xu, 2010; Kohut et al., 2010; Longo and Fabrizio, 2012). Recently, it was shown that phosphorylation of adenylate cyclase by Snf1 in *S. cerevisiae* reduces its activity and elevates stress response, yet such a connection in filamentous fungi or fungal plant pathogen was never made (Nicastro et al., 2015a). Here, both Snf1 and Snf2 are phosphorylated following MMS exposure (Fig. 8), therefore phosphorylation by Snf1 of adenylate cyclase is expected to contribute to activation of the MMS response in *F. oxysporum*. Nicastro et al. envisioned the Snf1 effects on the stress response to be mediated by PKA (Nicastro et al., 2015a; Nicastro et al., 2015b). However, adenylate cyclase partners with other proteins like Sgt1 that are phosphorylated in our database. It is not clear how adenylate cyclase phosphorylation affects non-PKA interactions.

While we clearly see a stress response and massive activation of various error- free DNA repair mechanisms (direct reversal, BER, NER, homologous recombination, template switching, Fig. 4, 5) it is unclear whether or not a mutagenesis program is activated. This is because the two major translesion DNA polymerases Rev3 and Rad30 are downregulated following MMS exposure, while others, like DNA polymerase kappa, are upregulated (Fig. 5). In any event, the amount of upregulation of mutagenic DNA polymerases is not even close to activation of error-free repair. Indeed, there is only three fold induction of mutation frequency following MMS exposure. In yeast, exposure to the same dose but only for half an hour resulted in similar induction of mutation but with no effect on survival (Ma et al., 2008; Yang et al., 2010).

### 4.4 Concluding remarks

While we point-out to many similarities in the MMS response of *S. cerevisiae* and *F. oxysporum* (Yap1, Xbp1, proteasome), the number of differentially expressed genes in *F. oxysporum* following MMS exposure is bigger than the number of genes in *S. cerevisiae* altogether. It boils down to the necessity to study in detail the role of filamentous-fungi specific or even *Fusarium*-specific genes in the response to MMS. Frequent genome rearrangements were observed in *F. oxysporum* and related fungi (Boehm et al., 1994; De Jonge et al., 2013; Shahi et al., 2016) our analysis points to the modules that guard the *Fusarium* genome understanding how these modules are activated or de-activated during the life cycle and of the pathogen is going to be a key in understanding its genome evolution.

## 5. Acknowledgements

This work was supported by Israel Science Foundation Grant 211/16 to S.C and BARD grant IS-4932-16 to SC and LJM.

---

**Fig. S1.**
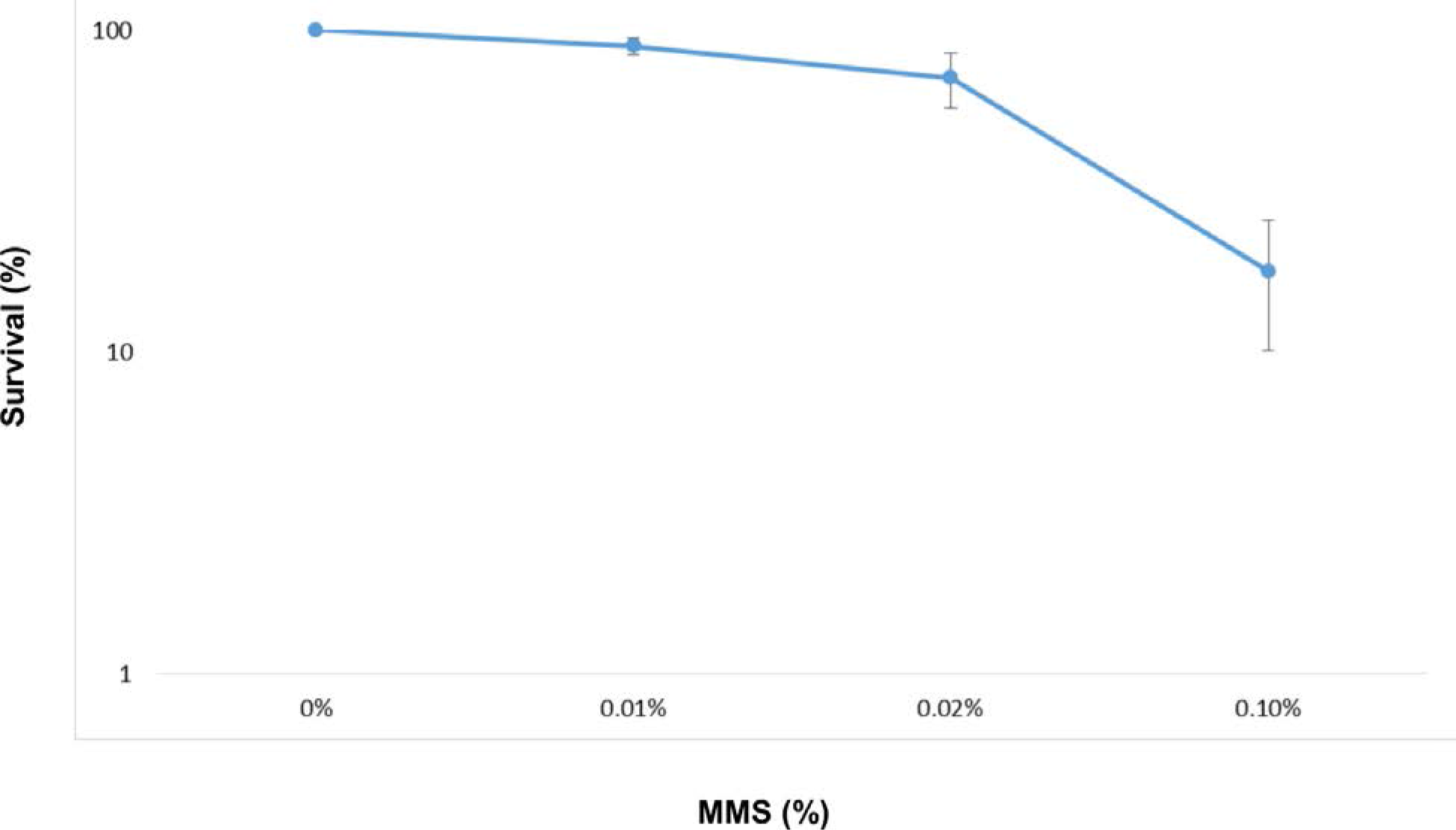
Dose response survival curve of MMS treated cells. Conidia were inoculated in PDB for five hours then treted with the indicated MMS doses for another three hours. Finally a sample of the culture was spread with appropriate dilutions on PDA plates (W/O MMS). Colonies were counted after three days, survival rate was calculated relatively to untreated cultures (eight hours inoculation in PDB).

**Fig. S2.**
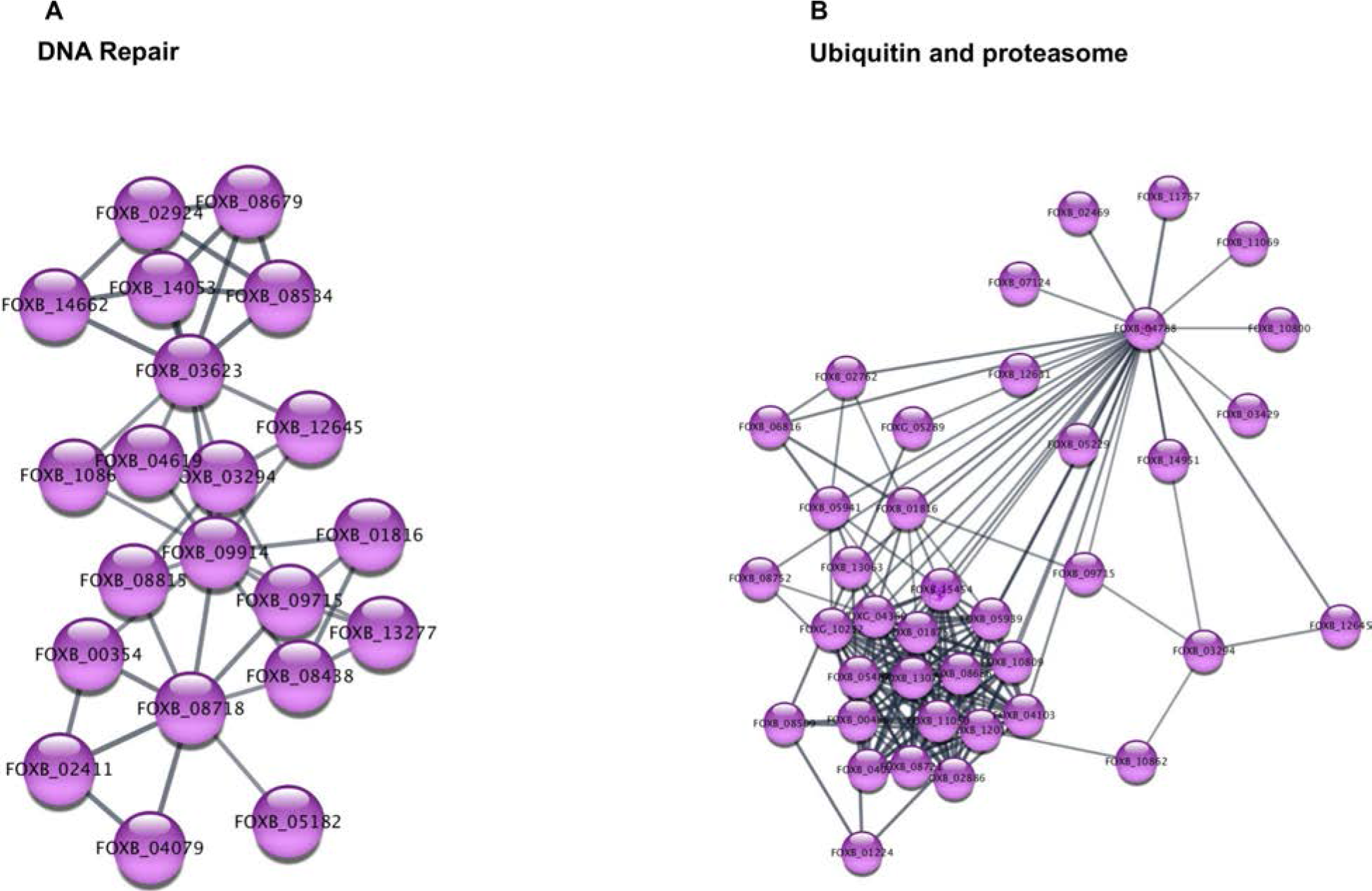
Cellular modules that are up-regulated in response to exposure of 0.05% MMS. Conidia was inoculated for five hours in PDB, then exposed to 0.05% MMS for another three hours. RNA was isolated and converted to RNA sequencing library using the 3’ quantseq kit of Lexogen (see details in text and under Materials and Methods). Up-regulated genes (fold increase 2 < and p < 0.05) were analyzed using STRING platform similarly to Fig. 2 with modifications; *Fusarium oxysporum* grid was used and confidence was set to be high (0.7). The number of nodes in the dataset was significantly higher than expected by random (the whole genome, p < 10^−16^). The biggest clusters of proteins are presented using cystoscope as a visualization tool.

**Fig. S3.**
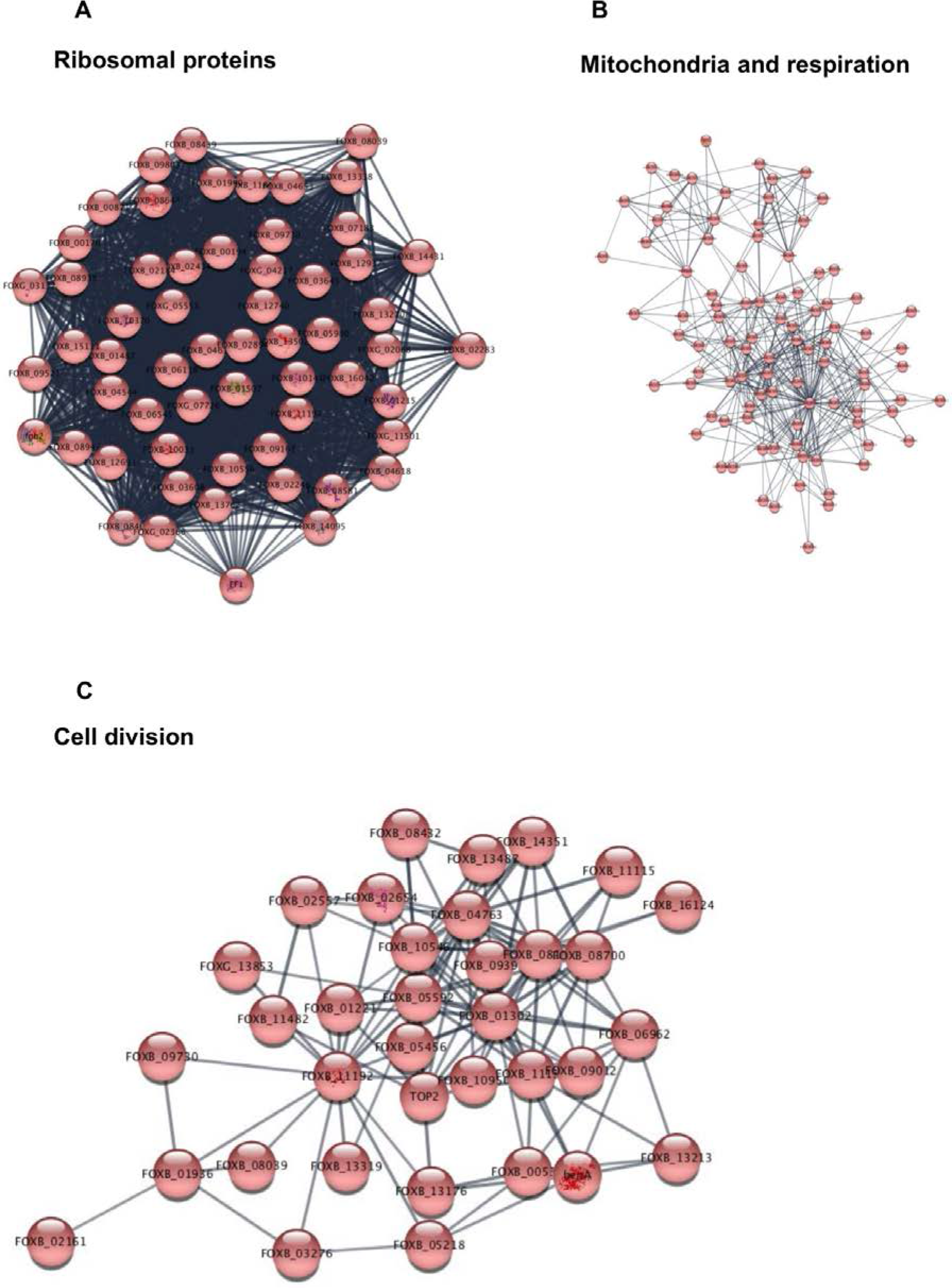
Cellular modules that are down-regulated in response to exposure of 0.05% MMS. As described for Fig. S2 but MMS down-regulated genes were analyzed.

**Fig. S4.**
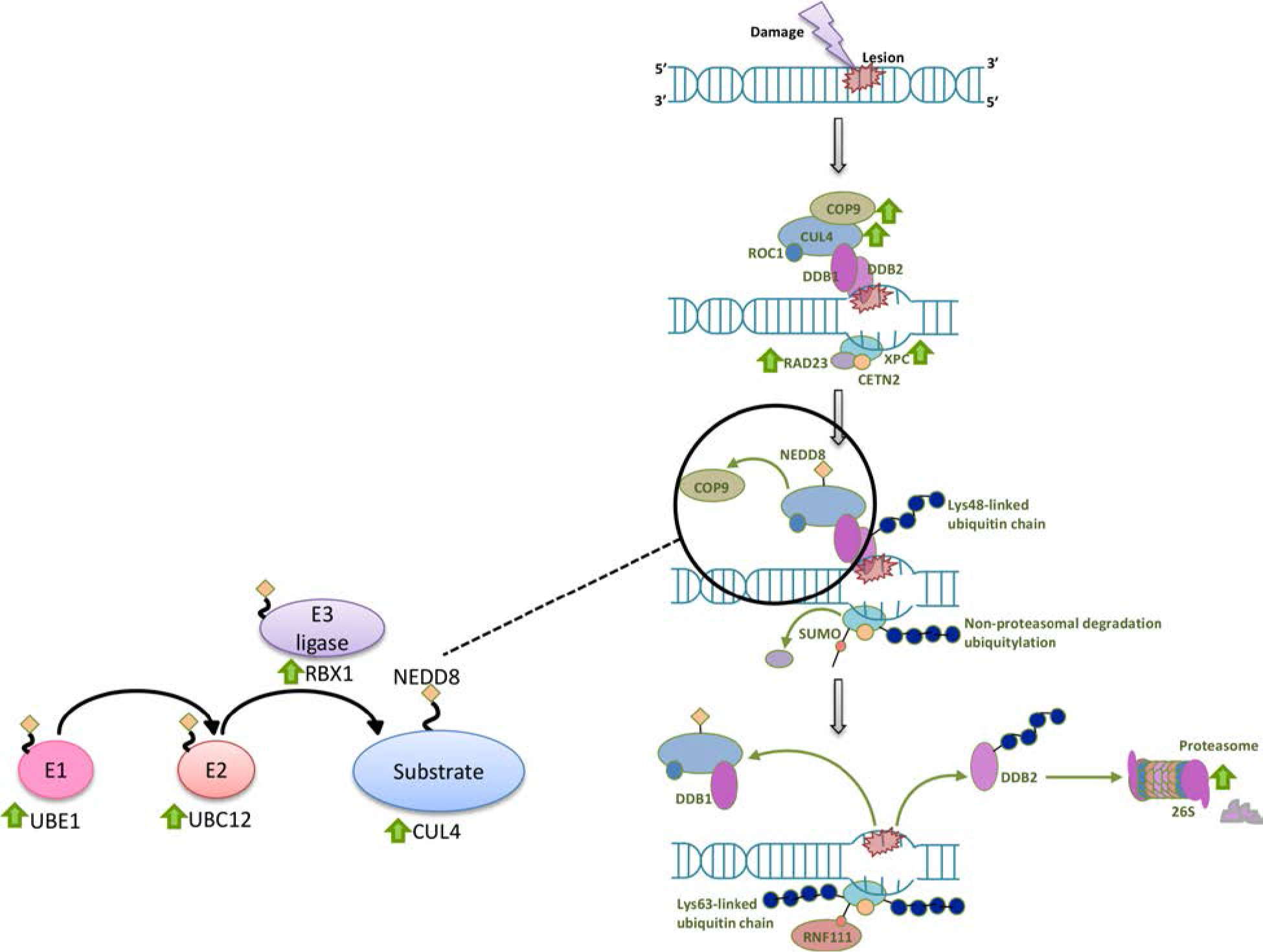
Nucleotide excision repair and Nedd8-Cul4-Cop9 pathways are activated following MMS exposure. A scheme of global nucleotide excision repair pathway is presented. Within the scheme, genes that are up-regulated following MMS exposure are indicated by a green arrow. Cop9-Nedd8-Cul4 proteins are presented in the major scheme and the detailed neddylation reaction is described at the accompanied scheme as a zoom-in of the encircled image.

**Fig. S5.**
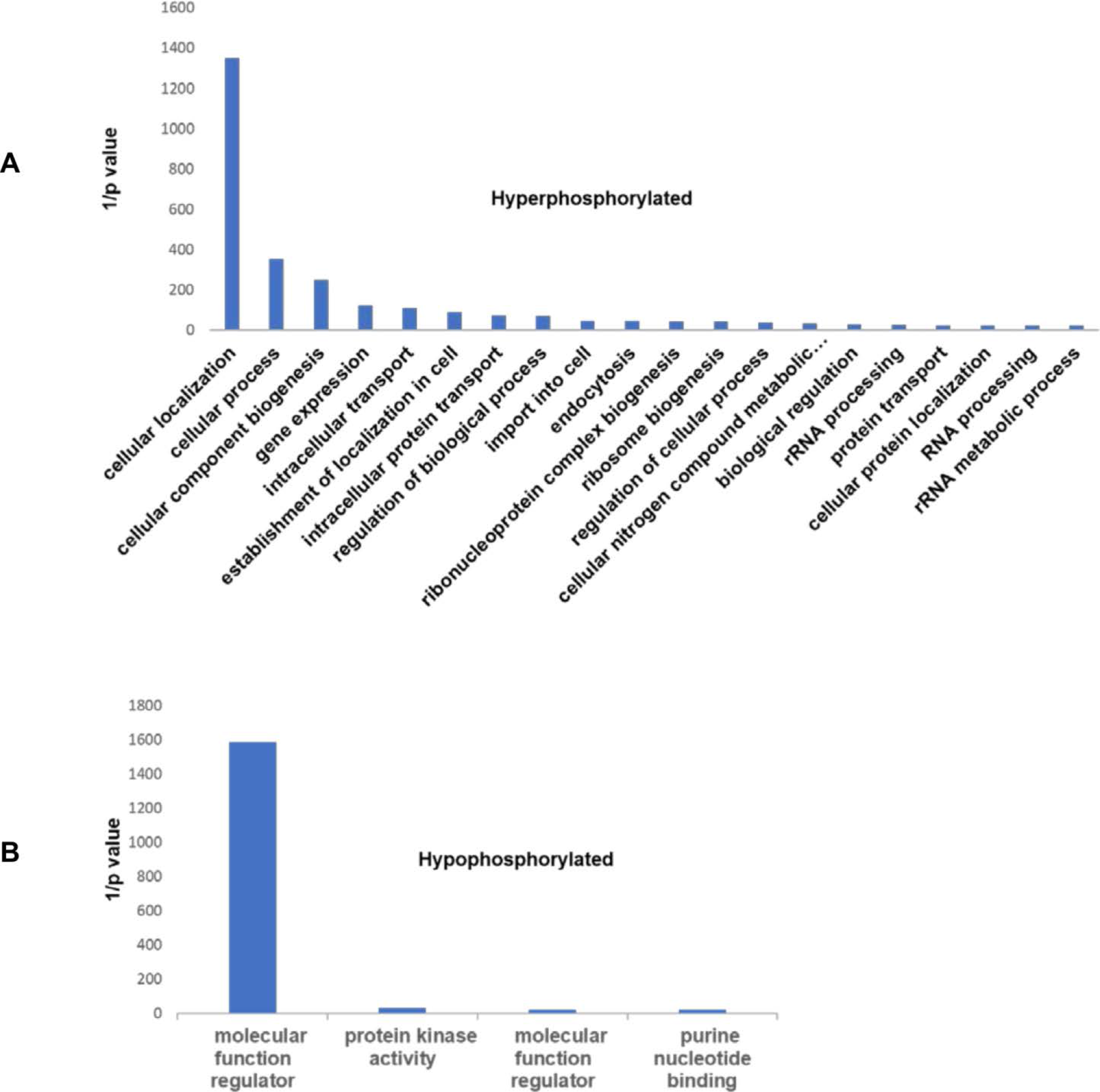
Enriched Gene Ontology terms among differentially phosphorylated proteins in *F. oxysporum* following MMS exposure. Classification of phosphorylated (A) and de-phosphorylated (B) proteins of *F. oxysporum* f.sp *lycopersici* exposed to 0.1% MMS for three hours to gene ontology (GO) terms. GO term finding was done as described under Materials and Methods; differentially expressed genes were identified in Table S8 (2 fold < and p < 0.05)

